# Nuclear receptor subfamily 4A signaling as a key disease pathway of CD1c+ dendritic cell dysregulation in systemic sclerosis

**DOI:** 10.1101/2021.11.08.467605

**Authors:** N.H. Servaas, S. Hiddingh, E. Chouri, C.G.K. Wichers, A.J. Affandi, A. Ottria, C.P.J. Bekker, M. Cossu, S. Silva-Cardoso, M. van der Kroef, A.C. Hinrichs, T. Carvalheiro, N. Vazirpanah, L. Beretta, M. Rossato, F. Bonte-Mineur, T.R.D.J. Radstake, J.J.W. Kuiper, M. Boes, A. Pandit

**Author notes:** These authors contributed equally (J.J.W. Kuiper, M. Boes, A. Pandit). Corresponding authors: Dr. Aridaman Pandit, Center for Translational Immunology, Wilhelmina Children’s Hospital, University Medical Centre Utrecht, Lundlaan 6, 3584 EA Utrecht, The Netherlands., Dr. Marianne Boes, Center for Translational Immunology, Wilhelmina Children’s Hospital, University Medical Centre Utrecht, Lundlaan 6, 3584 EA Utrecht, The Netherlands.

## Abstract

**Objectives:** To identify key disease pathways driving conventional dendritic cell (cDC) alterations in Systemic Sclerosis (SSc).

**Methods:** Transcriptomic profiling was performed on peripheral blood CD1c+ cDCs (cDC2s) isolated from 12 healthy donors and 48 SSc patients with all major disease subtypes. Differential expression analysis comparing the different SSc subtypes and healthy donors was performed to uncover genes dysregulated in SSc. To identify biologically relevant pathways, a gene co-expression network was built using Weighted Gene Correlation Network Analysis. We validated the role of key transcriptional regulators using ChIP-sequencing and *in vitro* functional assays.

**Results:** We identified 17 modules of co-expressed genes in cDC2s that correlated with SSc subtypes and key clinical traits including auto-antibodies, skin score, and occurrence of interstitial lung disease. A module of immune regulatory genes was markedly down regulated in patients with the diffuse SSc subtype characterized by severe fibrosis. Transcriptional regulatory network analysis performed on this module predicted *NR4A* (nuclear receptor 4A) subfamily (*NR4A1, NR4A2, NR4A3*) genes as the key transcriptional mediators of inflammation. Indeed, ChIP-sequencing analysis supported that these NR4A members target numerous differentially expressed genes in SSc cDC2s. Inclusion of NR4A receptor agonists in culture-based experiments provided functional proof that dysregulation of NR4As affects cytokine production by cDC2s and modulates downstream T-cell activation.

**Conclusions:** NR4A1, NR4A2 and NR4A3 are important regulators of immunosuppressive and fibrosis-associated pathways in SSc cDC2s. Thus, the NR4A family represent novel potential targets to restore cDC homeostasis in SSc.

**KEY MESSAGES:** *What is already known about this subject?:*

- CD1c+ conventional dendritic cells (cDC2s) are implicated as key players in Systemic Sclerosis (SSc), but key molecular mechanisms underlying their dysregulation were unknown.

*What does this study add?:*

- Transcriptomic analysis and network analysis identified modules of coexpressed genes in cDC2s that correlated with SSc subtypes and key clinical traits.
- The *NR4A* (nuclear receptor 4A) subfamily (*NR4A1, NR4A2, NR4A3*) genes act as master regulators of key immune regulatory genes dysregulated in SSc cDC2s, as shown by multi-omics integration analysis using transcriptomics and targeted ChIP-sequencing.
- Pharmacological activation of NR4As inhibits pro-inflammatory cytokine production and CD4+ T-cell activation by cDC2s.

*How might this impact on clinical practice or future developments?:*

- NR4As are attractive candidates for novel treatment options to attenuate pro-inflammatory and pro-fibrotic responses in SSc patients.

## INTRODUCTION

Systemic Sclerosis (SSc) is a complex, chronic autoimmune disease mainly characterized by vascular abnormalities, immunological abnormalities, and fibrosis of the skin and internal organs[1]. According to the American College of Rheumatology (ACR) criteria, and their extent of skin fibrosis, patients are classified into four subsets: early (eaSSc), non-cutanous (ncSSc), limited (lcSSc) and diffuse (dcSSc)[2], [3]. Vascular injury appears to be one of the earliest events in the pathogenesis of SSc[4], which in turn leads to the recruitment and activation of immune cells secreting pro-inflammatory cytokines and growth factors[5]. The resulting mix of inflammatory molecules induces the differentiation of resident epithelium, endothelium, monocytes and fibroblasts into myofibroblast that deposit excessive amounts of extracellular matrix, eventually leading to permanent tissue scarring[6], [7].

Conventional dendritic cells (cDCs) are a population of antigen presenting cells that play a central role in regulating adaptive immune cell responses[8], but also vascular tissue, and fibroblasts[9], [10]. Given their placement at the crossroads of inflammation and fibrosis, cDCs are implicated in the pathogenesis of SSc, and are hypothesized to be essential for the activation of pathways that promote fibrosis[11]. Indeed, in the early phases of the disease cDCs migrate to the skin [12], [13] and display an enhanced pro-inflammatory and pro-fibrotic cytokine production upon innate immune stimulation by Toll like receptor (TLR) ligation [14]. Furthermore, cDCs are a rather heterogeneous population, with CD1c+ cDCs (cDC2s) having a high capacity for priming CD4+ T cells[15]. Given the well-established pathogenic role of T cells in SSc[16]–[19], cDC2s are a highly interesting subset to investigate in the disease.

Although the data from previous studies supports an instructive role for cDCs in the pathogenesis of SSc, the molecular mechanisms that drive their dysregulation in the disease remain incompletely understood. To study the mechanisms behind cDC dysregulation in SSc, we compared the transcriptomic profile of circulating cDC2s obtained from peripheral blood (PB) of SSc patients and healthy controls. Using co-expression network analysis and *in vitro* validation studies, we aimed to unravel key players of cDC2 dysregulation in SSc.

## METHODS

PB samples were collected from patients with SSc and age/sex matched healthy controls (HC) from the University Medical Center Utrecht (The Netherlands), the Maasstad Medical Center Rotterdam (The Netherlands), and the IRCCS Policlinico of Milan (Italy) (Table 1). All participants signed an informed consent in adherence to the Declaration of Helsinki Principles. All patients fulfilled the ACR/EULAR 2013 classification criteria for SSc[2]. We also included in our studies ncSSc patients who fulfilled the classification criteria but did not present skin fibrosis, and eaSSc patients with Raynaud’s Phenomenon and positivity for SSc-specific autoantibodies and/or typical nailfold capillaroscopy patterns, as defined by LeRoy et al[3]. For functional experiments on healthy cDC2s not paired with SSc patients, we used buffy coats (Sanquin, Amsterdam, The Netherlands).

**Table 1.**
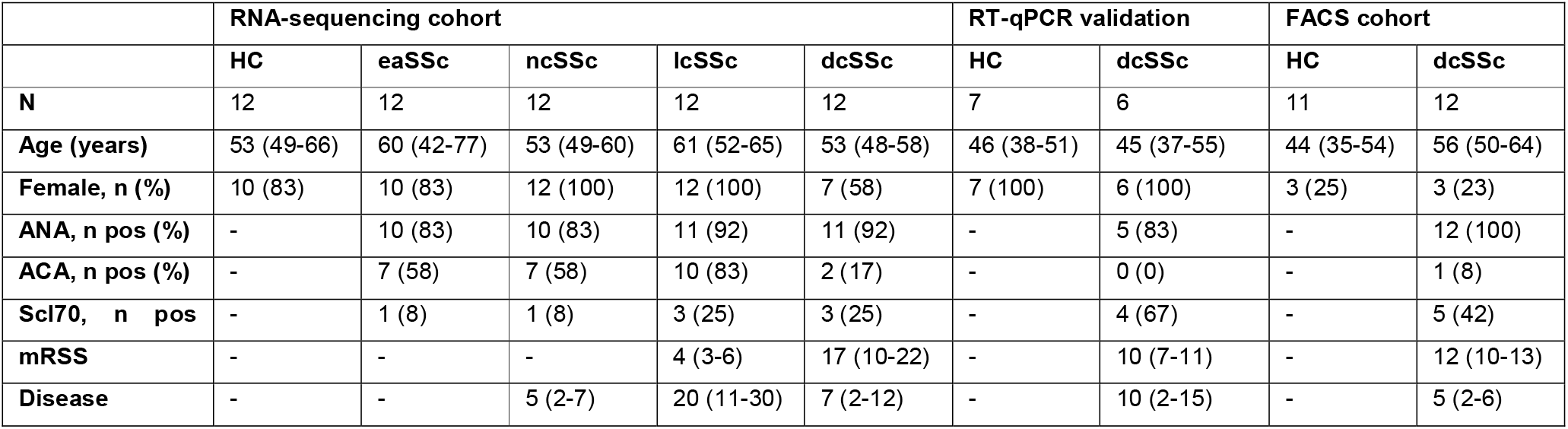
Clinical characteristics of subjects enrolled in the study. Reported values indicate the number (n) of individuals, with the (%) for each parameter. For age, mRSS and disease duration (in years), the median with interquartile range is given. ANA, anti-nuclear antibodies; ACA, anti-centromere antibodies; Scl70, antitopoisomerase antibodies; mRSS, modified Rodnan skin score; HC, healthy controls; eaSSc, early SSc; ncSSc, non-cutaneous SSc; lcSSc, limited cutaneous SSc; dcSSc, diffuse cutaneous SSc; pos, positivity.

For RNA-sequencing analysis, pair wise comparisons between patients groups and HC were made using the Wald test implemented in DESeq2, and genes with a nominal *P*<0.05 were considered to be significantly differential. Unless indicated otherwise, for functional experiments, the Mann–Whitney U test was used to compare any combination of two groups. For multiple group comparisons, the one- or two-way analysis of variance (ANOVA) was used. A *P*<0.05 was considered statistically significant. Detailed description of experimental methods used is available in the Supplemental Materials and Methods.

## RESULTS

### SSc cDC2s are transcriptionally distinct from healthy cDC2s

RNA sequencing of purified blood cDC2s from 48 SSc patients and 12 matched HCs was performed to assess differences in their transcriptomic profiles. We found 6,594 differentially expressed genes (DEGs) in at least one SSc subset compared to healthy controls (*P*<0.05) (**Figure 1A/B**). Principal component analysis (PCA) revealed that these genes generally separated SSc patients from HCs (**Figure 1C**). GO-term enrichment analysis showed that DEGs were predominantly enriched in pathways related to immune cell activation, interferon signaling and translation (**Figure 1D, Supplementary Table 1**). These results indicate that cDC2s from SSc patients have a distinct transcriptional profile from healthy cDC2s. Accordingly, the identified DEGs may allude to pathways relevant for SSc pathogenesis.

**Figure 1.**
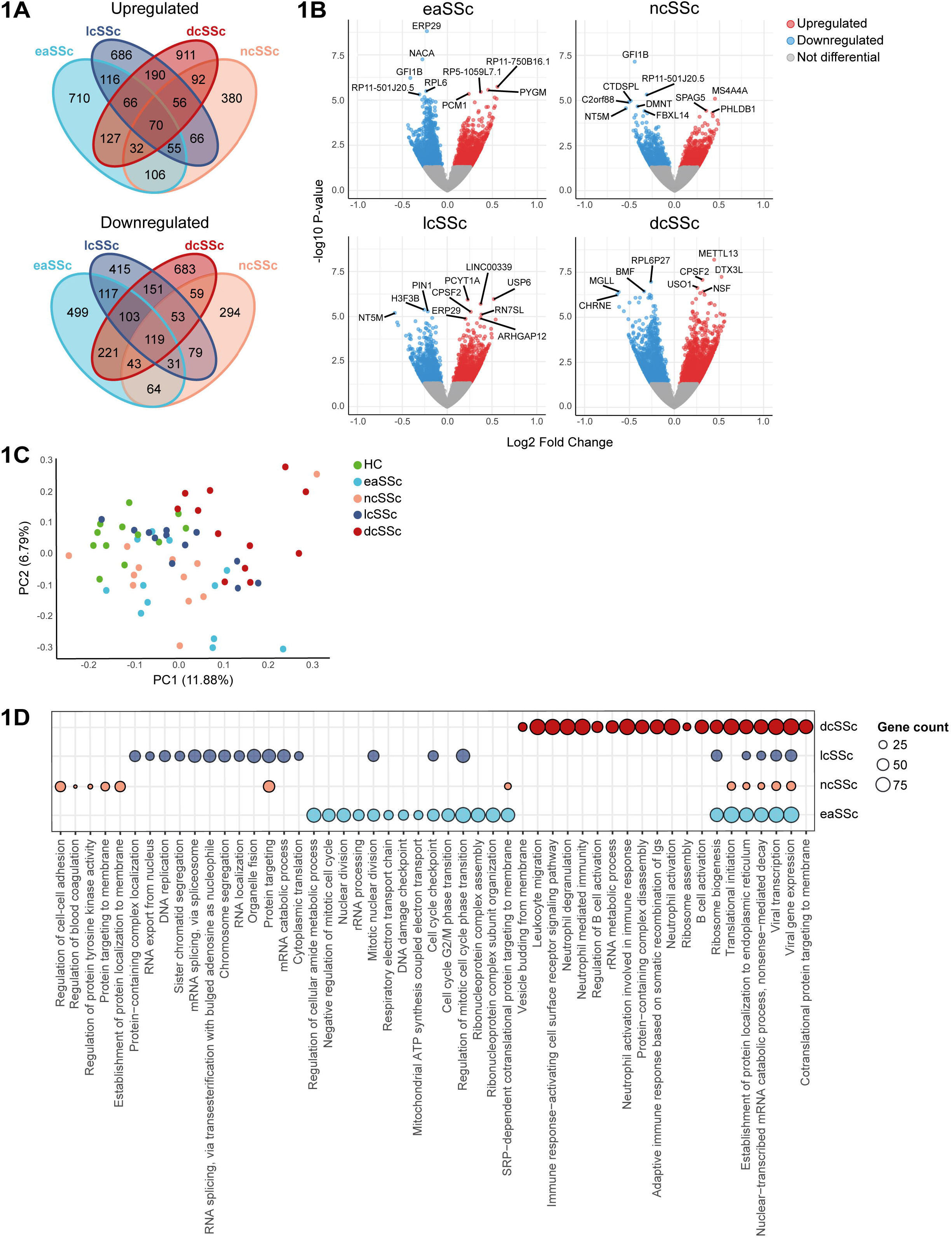
SSc cDC2s are transcriptomically distinct from healthy cDC2s. **A)** Number of differentially expressed genes (*P*<0.05) identified in cDC2s from different SSc subsets versus healthy controls. **B)** Volcano plots highlighting transcriptional differences between different SSc subsets and healthy controls. Blue dots represent significantly downregulated genes (*P*<0.05, log_2_ fold change < 0), while red dots represent significantly upregulated genes (*P*<0.05, log_2_ fold change > 0). Top differentially expressed genes based on p-value are highlighted. **C)** Principal component analysis (PCA) of the differentially expressed genes from all comparisons of SSc patients to healthy controls. **D)** Pathway enrichment analysis of differentially expressed genes identified in SSc patients versus healthy controls. Circle size denotes the number of differentially expressed genes associated to enriched pathways. When available. top 10 pathways are shown (B&H corrected *P* < 0.05).

### NR4As are key transcriptional regulators of functionally relevant pathways altered in SSc cDC2s

To further study the molecular pathways dysregulated in SSc cDC2s, we constructed a genome-wide gene co-expression network[20]. Following this approach we identified 42 modules of tightly co-expressed genes in cDC2s. From these, we identified 17 clinically relevant modules that were enriched in DEGs and also displayed a significant correlation with clinical traits relevant for SSc (**Figure 2A**). Functional annotation of these clinically relevant modules showed that some of them were associated with molecular pathways relevant in the context of cDC biology and inflammation. These included the blue module (associated with viral pathways and ribosomes), the darkgreen module (associated with immune cell activation) and the violet module (associated with antigen presentation and inflammation) (**Figure 2B**).

**Figure 2.**
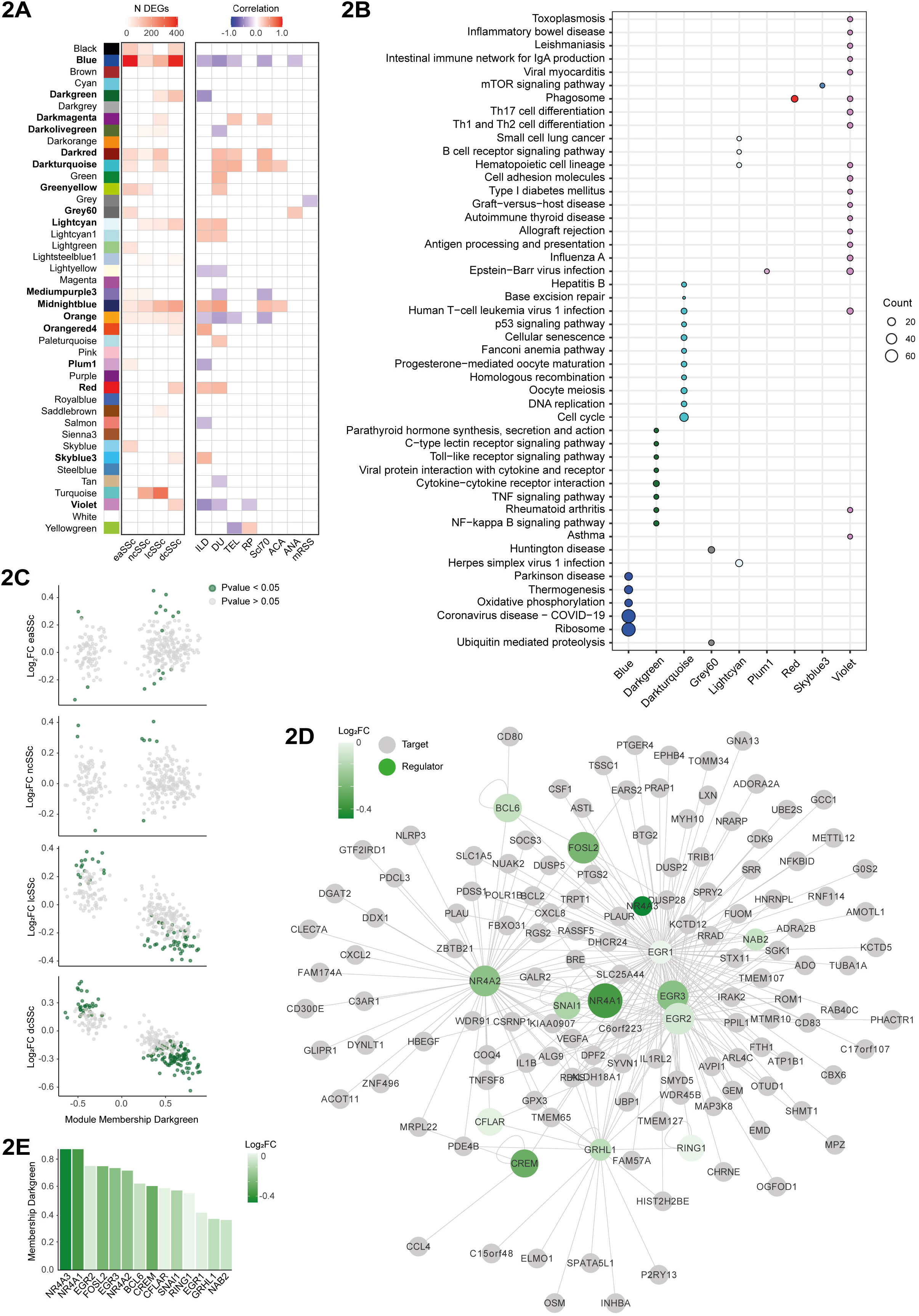
Co-expression network analysis identifies NR4A transcription factors of regulatory pathways decreased in SSc cDC2s. (**A**) Heatmaps depicting overlap of differentially expressed genes with co-expression modules (left) and correlation of module eigengenes (MEs) to SSc clinical traits (right). Cells of significant enrichments for differentially expressed genes (Fisher’s exact test *P*<0.05) or significant correlations with clinical traits (Pearson, *P*<0.05) are highlighted. N DEGs = number of differentially expressed genes. Modules significantly enriched in DEGs and correlated with clinical traits are highlighted in bold. ILD = interstitial lung disease, DU = digital ulcers, TEL = telangiectasia, RP = Raynaud’s phenomenon, Scl70 = anti-Scl-70 antibodies, ACA = anti-centromere antibodies, ANA = antinuclear antibodies, mRSS = modified rodnan skin score. (**B**) KEGG enrichment of relevant modules. Circle size denotes the number of module genes associated to enriched pathways. When available, top 10 pathways are shown (B&H corrected *P* < 0.05). (**C**) Module membership (x-axis) and the Log_2_FC (fold change) in gene expression compared to healthy controls (y-axis) for genes in the darkgreen module in patients with eaSSc, ncSSc, lcSSc and dcSSc. Significantly differentially expressed genes are highlighted in green (*P*<0.05). (**D**) Transcription factor network obtained for the darkgreen module. Transcriptional regulators (green) are connected to their targets (grey) based on known interactions from REGNET and TTRUST databases. Green shading indicates fold change between dcSSc and healthy cDC2s for transcriptional regulators. Node size denotes module membership in the darkgreen module. (**E**) Module membership (y-axis) and Log2FC (healthy versus dcSSc, colour scale) of transcriptional regulators (x-axis) identified in the darkgreen module.

Next, we focused on the darkgreen module given its association with immune cell activation. Closer inspection of this module revealed that genes with the highest module membership (MM; i.e., best representative of the overall module gene expression pattern), were also strongly downregulated in dcSSc patients (**Figure 2C**). This indicates that genes downregulated in dcSSc cDC2s are likely driving this module. Because transcription factors (TFs) are key regulators of gene expression, we next constructed a TF-target network based on known interactions from REGNET and TTRUST databases. Following this approach, we identified 13 TFs in the darkgreen module that had target genes present in the same module (**Figure 2D**). Most notable were members of the NR4A family of nuclear receptors, *NR4A1* and *NR4A2*, which have previously been described as important regulators of inflammation and fibrosis[21]–[26]. Of all TFs identified, *NR4A1* and *NR4A3* displayed the highest module membership (MM=0.87) and strongest downregulation in dcSSc patients (*NR4A3*: log2FC=-0.44 and *P*=0.0006; *NR4A1*: log2FC=-0.38 and *P*=0.004, **Figure 2E**). These, results support the importance of NR4As as regulatory factors in cDC2s, and suggests that their downregulated expression in dcSSc patients is of clinical relevance.

To validate the downregulation of *NR4As* in dcSSc patients (**Figure 3A**), we evaluated their expression by RT-qPCR in another, independent cohort of cDC2s from 6 dcSSc patients and 7 HCs (**Table 1**). *NR4A2* and *NR4A3* were significantly downregulated in dcSSc patients in this cohort (*P*=0.002 and *P*=0.035, **Figure 3B**), while for *NR4A1* we observed a trend for downregulation, although not significant (*P*=0.18, **Figure 3B**). Nonetheless, expression levels of all *NR4As* in the RNA-sequencing cohort and the RT-qPCR validation cohort were strongly correlated to each other (**Figure 3C**), with consistent downregulation of all NR4As in cDC2 of dcSSc patients.

**Figure 3.**
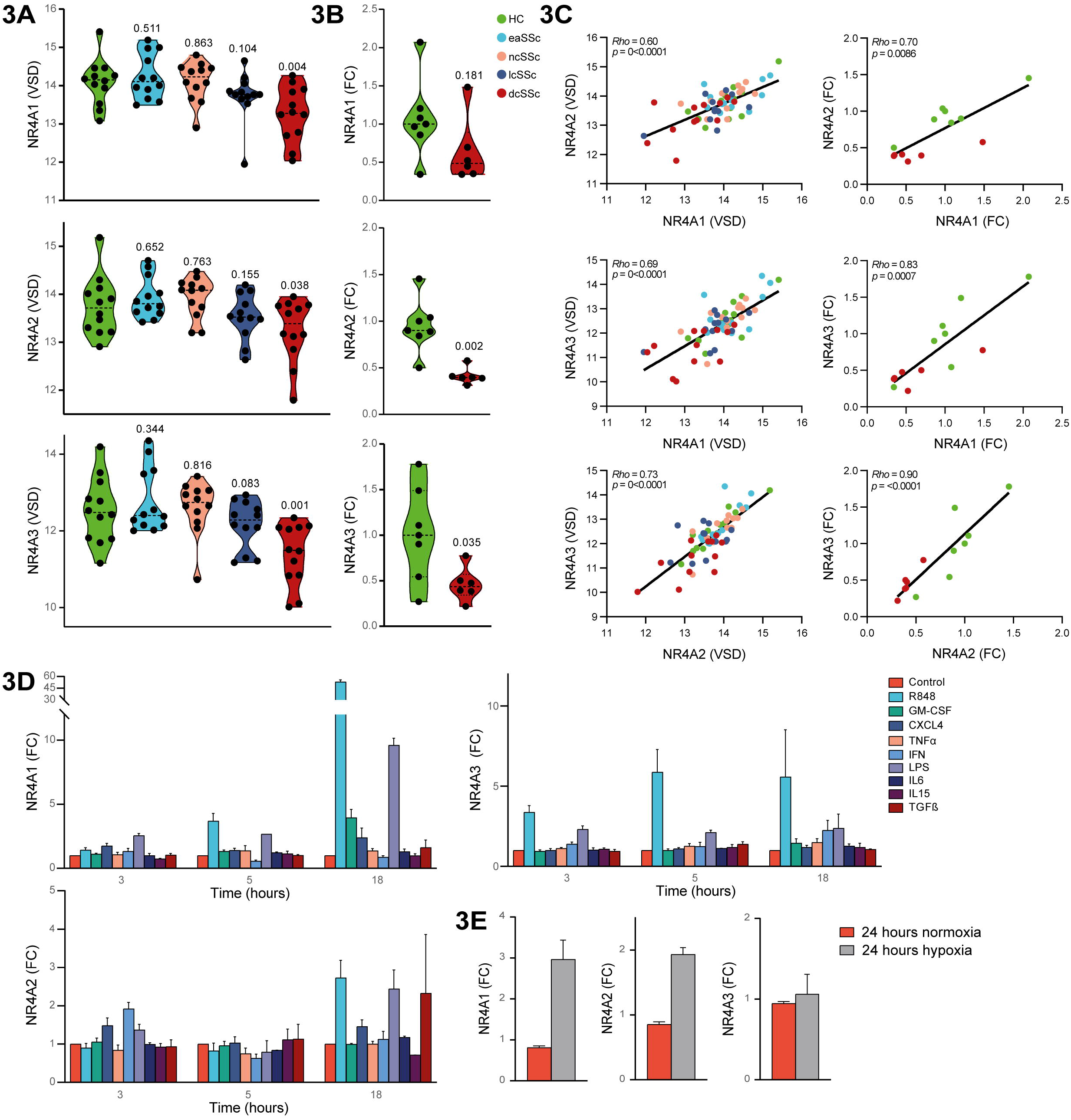
Characterization of NR4A expression in SSc and healthy cDC2s. (**A**) Violin plots depicting *NR4A* expression in SSc patients and healthy donors from the RNA-sequencing cohort (dashed lines indicates mean). For each comparison of SSc patients versus healthy controls, the p-value, calculated according to the Wald test implemented in DESeq2, is shown. (**B**) Violin plots showing *NR4A* expression in the validation cohort measured by target specific RT-qPCR. Data are reported as the fold change of each donor versus one representative healthy control. P-values (Kruskall–Wallis with post-hoc Dunn’s test), are reported. (**C**) Scatterplots showing the correlation (regression line) of *NR4A* expression in the RNA-sequencing cohort (left) and the validation cohort (right). Correlations were calculated using Spearman’s rank correlation coefficient (Rho). (**D**) *NR4A* expression in cDC2s from healthy controls stimulated for 3, 5 or 18 hours with Toll-like receptor ligands and cytokines implicated in dendritic cell biology and systemic sclerosis pathogenesis. Data are shown as mean with SEM of 3-4 experiments. (**E**) *NR4A* expression in cDC2s from healthy controls cultured in normoxic or hypoxic conditions. Data are shown as mean with SEM of 3 experiments.

### NR4A expression is induced by pro-inflammatory stimuli, yet downregulated in cDC2s obtained from dcSSc patients

To investigate whether activation of cDC2s from SSc patients associates with *NR4A* downregulation, we cultured freshly isolated cDC2s obtained from PB of healthy controls in the presence of various pro-inflammatory mediators for 3, 5, or 18 hours. *NR4A* expression was either induced or unaffected in all culture conditions studied (**Figure 3D**), with the strongest induction of expression observed upon stimulation with the TLR7/8 agonist R848 (Resiquimod) for 18 hours (**Figure 3D**). Additionally, since hypoxia has been described as an important inflammation related hallmark in SSc[27], we investigated the effect of hypoxia on *NR4A* expression in cDC2s. Expression of *NR4A1* and *NR4A2* was also induced in hypoxic conditions (**Figure 3E**). Next, to see whether cDC2s from dcSSc patients had a similar capacity as healthy cDC2s for activation-induced NR4A expression, we repeated the stimulation experiments on cDC2s obtained from 5 dcSSc patients and 5 matched HCs. No significant difference in *NR4A* expression was observed between R848 stimulated cDC2s from HC and SSc patients (**Supplementary Figure 1**). These results support that enhanced activation status of cDC2s does not underlie the decreased expression of *NR4As* observed in dcSSc patients, but is rather a result of an intrinsic mechanism in cDC2s.

### NR4As directly regulate crucial pathways dysregulated in cDC2s from SSc patients

To scrutinize the regulatory function of NR4As in cDC2s in steady state and during activation, we performed ChIP-sequencing for NR4A1, NR4A2, and NR4A3 in resting and stimulated cDC2s. We identified numerous NR4A binding sites, of which some were specific to resting conditions (medium), while others were specific to inflammatory conditions (R848) (**Supplementary Table 2**, **Supplementary Figure 2**). In line with their known roles in neuronal development[28] and cardiac tissue development[29], we found a significant enrichment of NR4A binding sites at genes related to these processes (**Figure 4A**). Moreover, we identified NR4A binding in the promoters of genes related to cell morphogenesis, extracellular matrix (ECM) organization and chemotaxis (**Figure 4A**). These results demonstrate that NR4As are involved in the direct transcriptional regulation of various key SSc-pathology related processes in cDC2s.

**Figure 4.**
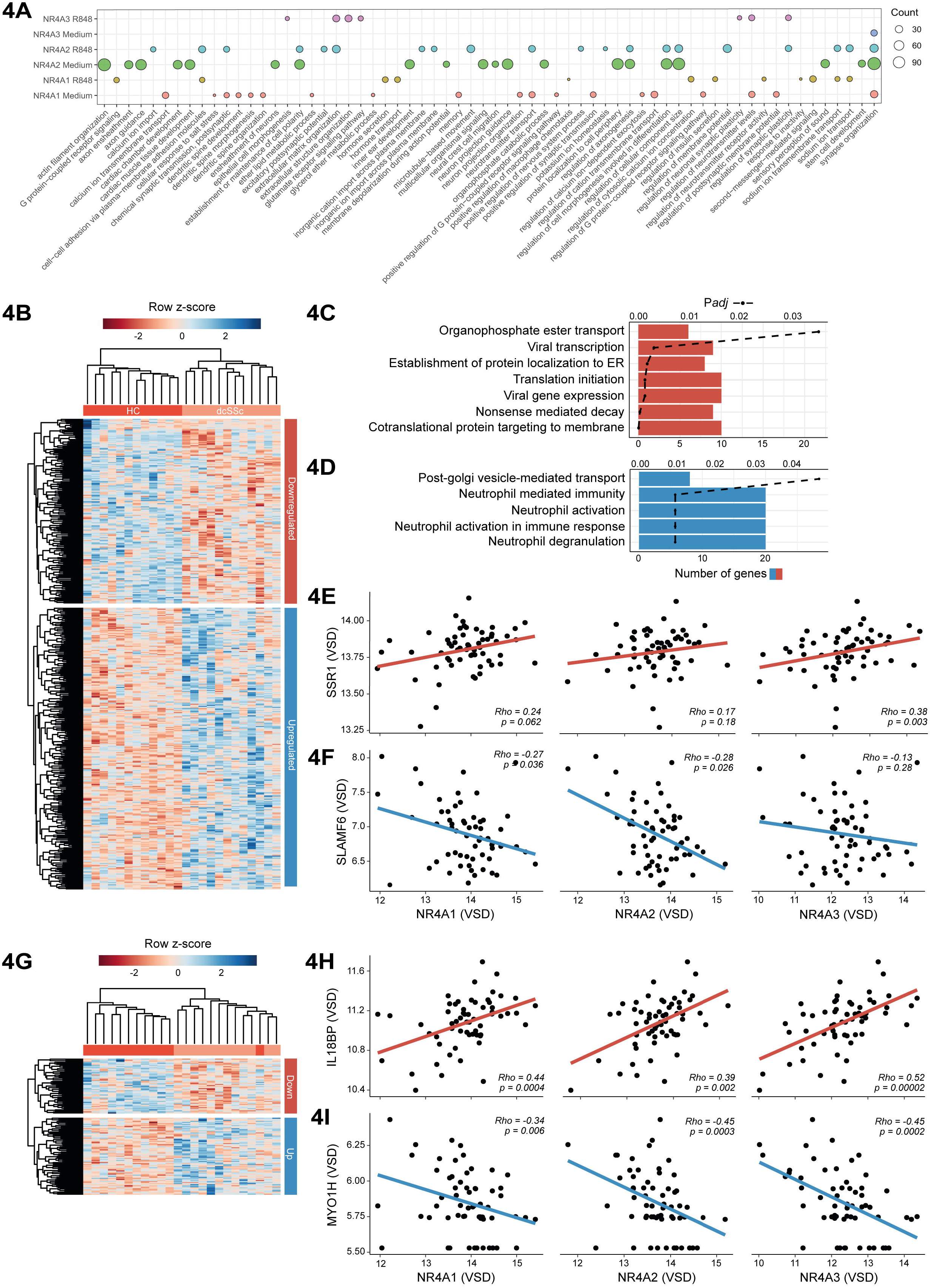
ChIP-sequencing of transcriptional regulation of cDC2s by NR4As. (**A**) GO-term enrichment of genes that show NR4A binding within their promoter region. Circle size denotes the number of genes associated to enriched biological processes. Top 20 are shown (B&H corrected *P*< 0.05). (**B**) Heatmap of differentially expressed genes in dcSSc that also display binding of NR4As at their promoters in resting cultured cDC2s. (**C**) Pathway enrichment of genes downregulated and (**D**) upregulated in dcSSc with NR4A binding at their gene promoters in resting cultured cDC2s. Terms for significantly enriched biological processes are given on the y-axis. Bars depict the number of genes identified within the enriched pathway (N genes, bottom x-axis), dashed line indicates B&H corrected p-value of the enrichment (p-value, top x-axis). (**E**) Scatterplots showing the correlation of *NR4A1-3* expression (x-axis) with *SSR1*, or (**F**) *SLAMF6* (y-axis). Correlations were calculated using Spearman’s rank correlation coefficient (Rho). VSD = variance stabilized data. (**G**) Heatmap of differentially expressed genes in dcSSc that also display binding of NR4As at their promoters in R848 stimulated cDC2s. (**H**) Scatterplots showing the correlation of *NR4A1-3* expression (x-axis) with *IL18BP*, or (**I**) *MYOH1* (y-axis). The red line on the scatter plots represents the regression line or “line of best fit” for positive correlations, and blue for negative correlations.

Many genes directly bound by NR4As in resting cDC2s in the ChIP-seq analysis were also differentially expressed in cDC2s from dcSSc patients in the RNA-seq analysis (**Figure 4B**). Moreover, in the RNA-seq cohort *NR4A* expression was correlated with genes associated with ER peptide translocation, including *SSR1* [30] (**Figure 4E**), and inversely correlated with genes related to immune activation, including *SLAMF6*[31] (**Figure 4F**). Additionally, we observed a strikingly lower overlap of DEGs with NR4A binding sites obtained from R848 stimulated cDC2s as compared to resting cDC2s (**Figure 4G**). This observation reflects a potential loss of NR4A binding during cDC2 activation in SSc. Among the genes directly bound by NR4As in stimulated cDC2s, we identified the anti-inflammatory IL18 binding protein (*IL18BP*)[32], which was downregulated in dcSSc cDC2s and displayed a direct correlation with *NR4As* in the RNA-sequencing cohort (**Figure 4H**). Among the upregulated genes we found Myosin IH (*MYO1H*), a molecule involved in cytokinesis, maintenance of cell shape, and cell motility[33], which was inversely correlated with *NR4A* expression (**Figure 4I**). Taken together, these results demonstrate that NR4As are directly involved in transcriptional programs limiting cDC2 activation *in vitro*, at least in the culture-based experimental conditions tested here. Accordingly, the loss of NR4A expression in SSc cDC2s leads to an increased activation of these cells.

### NR4A activation limits pro-inflammatory cytokine production by cDC2s from healthy controls and dcSSc patients

Next, we sought to identify whether activation of NR4A signaling could attenuate pro-inflammatory cytokine production by healthy and dcSSc cDC2s. We treated freshly isolated cDC2s with increasing concentrations of C-DIM5 (NR4A1 agonist) or C-DIM12 (NR4A2 agonist) and measured the expression *GUSB* to evaluate the effect of NR4A activation on viability. For C-DIM5 and C-DIM12, concentrations up to 10uM and 25uM were well tolerated (**Figure 5A**). Preincubation of cDC2s with 10uM C-DIM5 and C-DIM12 before stimulation with R848 led to a significant decrease in IL-6 production, both on mRNA and protein level (**Figure 5B**).These data confirmed the suppressive role of NR4As in pro-inflammatory cytokine production in cDC2s. To substantiate these results, and to evaluate the effect of NR4A agonists on dcSSc cDC2s, we repeated this experiment using PBMC samples obtained from 12 dcSSc patients and 11 matched HCs. Intracellular flow cytometry and gating on cDC2s (**Supplementary Figure 3**) again showed that NR4A activation led to a significant decrease of IL-6 production by these cells (**Figure 5C**). Notably, NR4A activation also led to a significant decrease of IL-6 production in dcSSc cDC2s, demonstrating that NR4A activation can effectively attenuate pro-inflammatory cytokine production in these patients.

**Figure 5.**
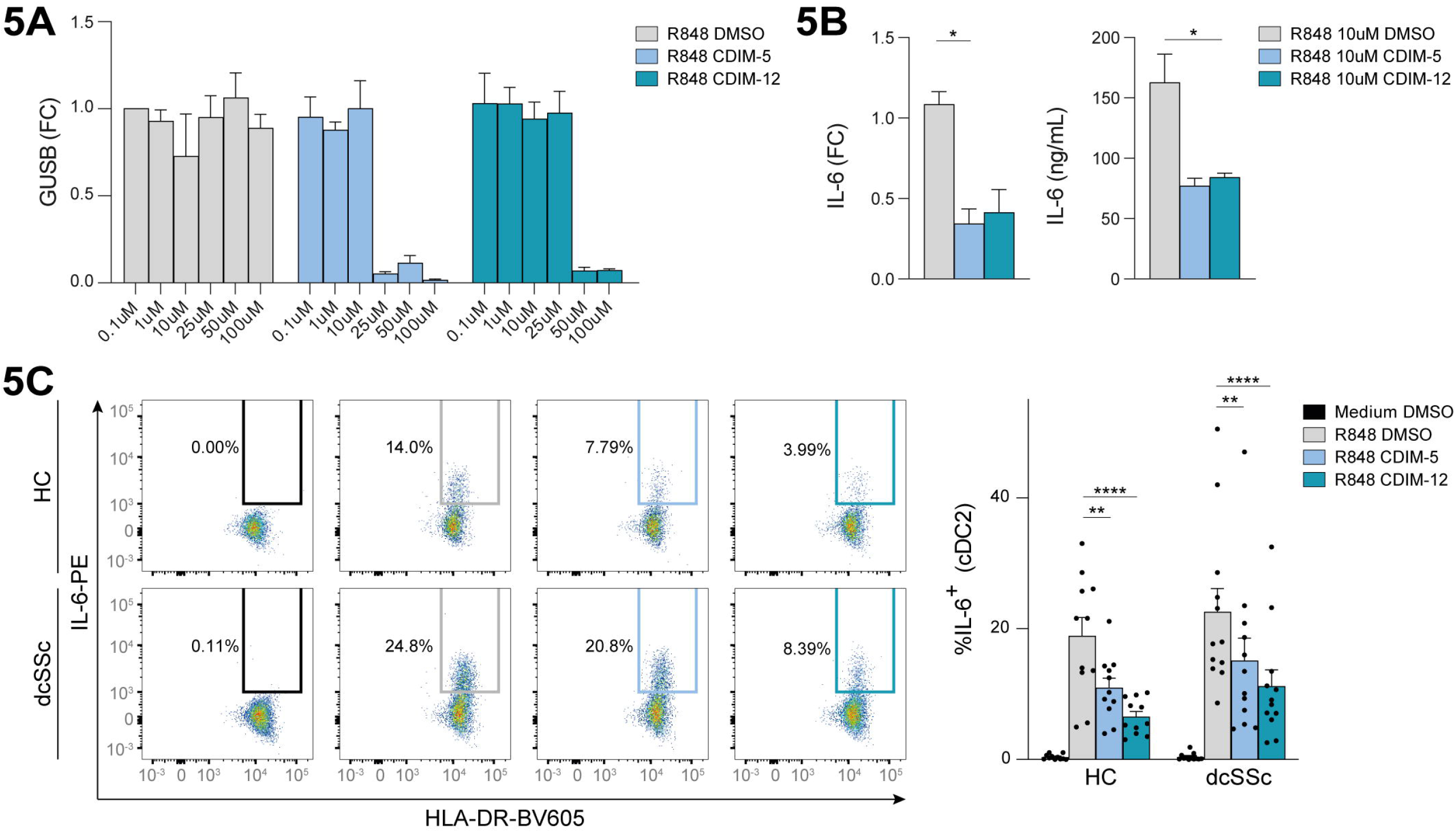
NR4A activation leads to a decrease in the production of IL-6 in healthy and dcSSc cDC2s. (**A**) RT-qPCR of *GUSB* mRNA expression by freshly isolated cDC2s after pre-incubation of increasing concentrations of DMSO (negative control), or NR4A agonists C-DIM5 and C-DIM12, followed by overnight stimulation with R848. Fold change (FC) in comparison to 1μM DMSO is shown. Data are shown as mean with SEM of 3 experiments. (**B**) *IL-6* mRNA (left) and protein expression (right) of cDC2s pretreated with 10uM DMSO, C-DIM5 or C-DIM12, followed by overnight stimulation with R848. Relative mRNA expression levels (FC) shown are normalized to *GUSB* housekeeping levels. Data are shown as mean with SEM of 3 experiments. * = *P* <0.05, calculated by one-way Anova followed by Friedman test for multiple comparisons. (**C**) Representative flow cytometry plots of the percentage of IL-6 positive cells within the cDC2 fraction in PBMC cultures pretreated with 10uM DMSO, C-DIM5 or C-DIM12, followed by overnight stimulation with R848. Barplots (mean + SEM) depict the quantification of flow cytometry data obtained from 12 healthy controls (HC) and 13 dcSSc patients. ** = *P* <0.01, **** = *P* <0.001, calculated by two-way Anova followed by Dunnett’s test for multiple comparisons.

### NR4A activation can decrease the CD4+ T-cell stimulatory capacity of cDC2s

Since cDC2s are indispensable for CD4+ T-cell activation, and CD4+ T-cells contribute to SSc pathogenesis[16]–[19], we next investigated the role of NR4As in priming cDC2s for autologous CD4+ T-cell activation (**Figure 6A, Supplementary figure 4**). cDC2s pretreated with NR4A agonists C-DIM5 and C-DIM12 were less capable of inducing IFNγ production by CD4+ T-cells compared to control (**Figure 6B**). These results held true for cDC2s stimulated with R848, showing that NR4A activation also attenuates CD4+ T-cell induction by cDC2s under pro-inflammatory conditions (**Figure 6C**). These data demonstrate that, beside controlling the expression of pro-inflammatory cytokines, NR4As also have the capacity to control T-cell activation by cDC2s.

**Figure 6.**
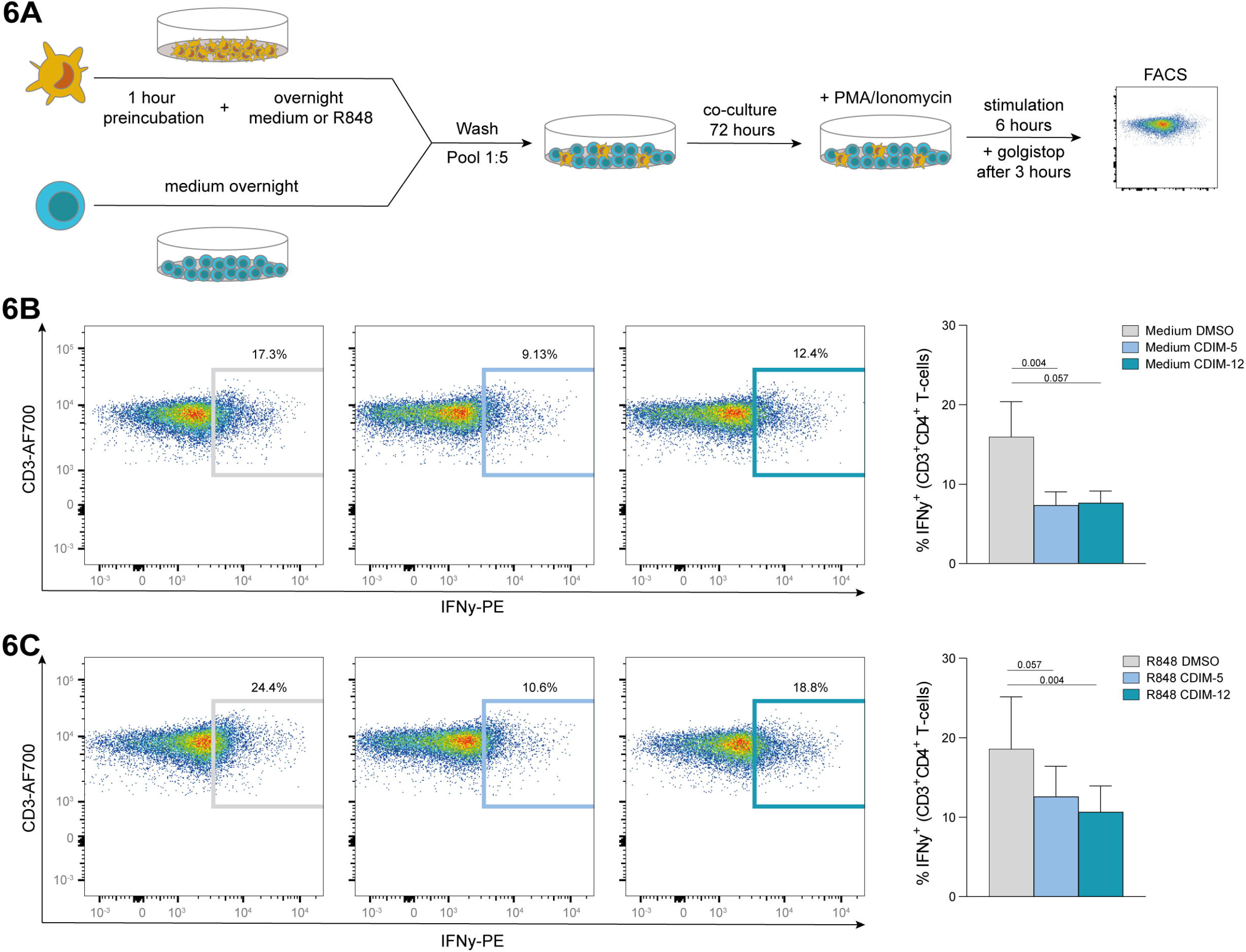
NR4A activation leads to a decrease in CD4+ T-cell activation by cDC2s. (**A**) Schematic overview of the co-culture setup of freshly isolated autologous CD4+ T-cells and cDC2s. (**B**) Representative flow cytometry plots of the percentage of IFNγ positive T-cells within the CD4+ T-cell fraction after 3 days of coculture with cDC2s pretreated with 10uM DMSO, C-DIM5 or C-DIM12, followed by overnight culture in medium or (**C**) overnight stimulation with R848. Barplots (mean + SEM) depict the quantification of flow cytometry data obtained from 5 donors. P-value in comparison to DMSO treatment is shown, calculated by one-way ANOVA followed by Friedman test for multiple comparisons.

## DISCUSSION

cDCs have been put forth as important immune cells involved in SSc pathogenesis[12]–[14]. However, key molecular mechanisms underlying their dysregulation were thus far unknown. Here we provide detailed transcriptomic profiling of cDC2s from SSc patients and HCs, which allowed us to characterize their transcriptomic landscape. We identified various clinically relevant modules of tightly co-expressed genes enriched in genes from pathways highly relevant for SSc and cDC biology, including immune cell activation, antigen presentation, anti-viral mechanisms, and translation. In particular, a module of immune regulatory genes was downregulated in cDC2s of dcSSc patients.TF network analysis pointed towards the NR4A family of orphan nuclear receptors as potentially important regulators of this module. These results implicate NR4As as key regulators of inflammatory responses by cDC2s in SSc.

NR4As regulate gene expression in a ligand-independent manner, meaning that their activity is largely dependent on their expression levels and posttranslational modifications[34], [35]. They play critical roles in the regulation of immune cell activation[36]–[38], and NR4A1 in particular, has been established as regulator of pro-inflammatory responses in DCs[24]. Our ChIP-seq analysis provides detailed insights into the direct binding of NR4As at their target gene promoters in primary human cDC2s. As ChIP-seq studies from these cells are very rare and mostly limited to model systems like monocyte derived DCs, these data provide a valuable resource for future studies on NR4As in general, as well as for cDC2s and SSc. We show that NR4As are strongly involved in transcriptional programs underlying DC dysregulation in SSc. Besides inflammation, these include the regulation of morphology and ECM production under stimulated conditions. Given the reduced expression of NR4As in circulating cDC2s from dcSSc patients, these results suggest that dcSSc cDC2s might show an enhanced expression of ECM related genes once they get activated, for example upon migration to the skin. Interestingly, inflammatory DCs have been implicated fibrosis in SSc through the increased secretion of the ECM molecules like fibronectin and α-SMA, thereby promoting myofibroblast differentiation[39]. Although more detailed ChIP-seq analyses to quantify NR4A binding in dcSSc in comparison to healthy donor cDC2s are needed to validate these results, our analysis points towards NR4As as major transcriptional regulators of pathways implicated in cDC dysregulation in SSc.

We show that activation of NR4As in cDC2s by selective agonists attenuates release of the pro-inflammatory cytokine IL-6 and downstream activation of CD4+ T-cells. Importantly, we show that, although the expression of *NR4A1-3* is downregulated in cDC2s obtained from dcSSc patients, activation of NR4A signaling by NR4A agonists C-DIM5 and C-DIM12 inhibits IL-6 production by cDC2s from these patients. Interestingly, IL-6 is an important cytokine linked to SSc pathogenesis, and levels of this cytokine are linked to worsening of clinical outcome and increased fibrosis[40]. Additionally, the NR4A1 agonist cytosporin-B has previously been shown to ameliorate collagen deposition and myofibroblast differentiation in bleomycin-induced mouse models of fibrosis[25], highlighting a role for NR4As at the cross-roads of fibrosis and inflammation. Although it remains to be investigated to what extend these anti-fibrotic effects are mediated through modulation of cDC2s, small molecule agonists that overcome the reduced expression of NR4A in cDC2s may represent potential targets for immunotherapy in SSc. Given the fact that SSc patients with early diffuse phenotypes have also previously been shown to display sign of enhanced DC activation with increased IL-6 production[14], targeting NR4As early in SSc pathogenesis might prevent DC activation at pre-fibrotic stages and limit disease progression.

Although factors that underlie the downregulated expression of NR4As in dcSSc cDC2s remain to be resolved, our experiments do provide new insights. In line with the roles of NR4As as immediate early response genes[34], stimulation of cDC2s with TLR ligands R848 and LPS, as well as hypoxia (linked to SSc pathogenesis[27]), did overall induce the expression of *NR4A1-3*. Also, cytokines known to be increased in peripheral blood of SSc patients or related to SSc pathogenesis, including CXCL4, IFNα, TGFβ, GM-CSF, IL-6 and IL-15 did not reduce NR4A expression, at least for the concentrations and time points included here. Moreover, stimulation of freshly isolated cDC2s from dcSSc patients with TLR7/8 ligand R848 also led to an induction of NR4A expression, comparable to healthy levels, suggesting that the upstream transcriptional regulation of NR4As is not defective. Given the well described heterogeneity of the cDC2 subset[41], [42], one might propose that the downregulation of NR4A expression that we observed in bulk cDC2s from dcSSc patients may reflect a disbalance of distinct populations within the cDC2 subset. Indeed expression of *NR4A2* and *NR4A3* has previously been shown to be low in CD1c^+^Tbet^-^ cDCs (also known as cDC2B), an inflammatory DC population within the cDC2 compartment[43]. However, recent analysis of the composition of cDC2 subsets in the blood of SSc patients by Dutertre *et al*., showed that the proportions of distinct cDC2 subpopulations in SSc was not different from the healthy situation[42]. Thus, the downregulation of NR4As that we observed in dcSSc cDC2s may not be attributed to heterogeneity within the DC compartment. It remains to be investigated what exact molecular mechanisms cause NR4A downregulation in dcSSc cDC2s. These might include alterations in the chromatin landscape or regulation at the post-transcriptional level.

In conclusion, we show that the NR4A transcription factor family members NR4A1, NR4A2 and NR4A3 are important regulators underlying cDC2 dysregulation in SSc. We propose that pharmacological activation of NR4As is an attractive candidate therapeutic option to attenuate pro-inflammatory and pro-fibrotic responses in SSc patients, in an untargeted manner, or using DC-directed approaches.

## ACKNOWLEDGEMENTS

We thank A. Pinheiro Lopes for her experimental advice on functional experiments with cDCs. Furthermore, we thank E. Ton, J.M. van Laar and J. Spierings from the Department of Rheumatology & Clinical Immunology in University Medical Centre Utrecht for their help with patient inclusion. We are grateful to the patients and healthy control donors who participated in this study.

## ETHICS APPROVAL

The study was approved by the medical ethics committee of the University Medical Center Utrecht (METC no. 12/296 and no. 13/697). All patients gave written informed consent in accordance with the declaration of Helsinki.

## DATA AVAILABILITY STATEMENT

Processed counts and raw files for RNA-sequencing data and ChIP-sequencing presented in this study have been deposited in NCBI’s Gene Expression Omnibus (GEO) database under GSE186199.

