## Supplementary Materials and Methods for "Nuclear receptor subfamily 4A signaling as a key disease pathway of CD1c+ dendritic cell dysregulation in systemic sclerosis"

**cDC2 purification**

Peripheral blood mononuclear cells (PBMCs) were isolated from whole heparinized-blood samples obtained from SSc patients and healthy volunteers, by density gradient centrifugation on Ficoll-PaqueTM Plus (GE Healthcare Life Sciences). The CD1c+ cDC population was purified from PBMCs using the MACS human CD1c (CD1c+) dendritic cell isolation kit (Miltenyi Biotec) on the autoMACS Pro Separator (Miltenyi Biotec) according to the manufacturer’s protocol. For the RNA-sequencing cohort as well as the RT-qPCR validation cohort, freshly isolated cDC2s were immediately lysed in RLTplus buffer (Qiagen) supplemented with β-mercaptoethanol, and stored at -20°C until further processing.

**RNA-sequencing and analysis**

Total RNA was purified from RLTplus lysates using the DNA/RNA/miRNA Universal Kit (Qiagen), according to manufacturer’s instructions. Purified RNA was quantified with the Qubit® RNA Assay Kit (Life Technologies) on the Qubit® Fluorometer (Invitrogen). RNA-sequencing was performed at the Beijing Genomics Institute (BGI). cDNA libraries were generated from total RNA using the TruSeq RNA sample preparation kit (Illumina), specifically selecting for polyadenylated transcripts. The libraries were sequenced on the HiSeq 2000 system (Illumina), using 100bp paired-end reads.

We obtained at least 20 million raw reads for each sample. After quality filtering according to the BGI pipeline, reads were aligned to the GrCh38 reference human genome and the homo sapiens transcriptome (Ensembl, version 79), using the STAR aligner[1]. Summed exon read counts per gene were calculated using the Python package HTSeq[2], using annotations from the GrCh38 built from the human genome (http://www.ensembl.org, version 79). Between lane normalization using the upper quartile normalization method was performed using the Bioconductor/R package EDAseq[3]. To account for batch effects arising from the inclusion of samples from different geographic locations (The Netherlands and Italy), we applied the generalized linear model from the Bioconductor/R package RUVseq[4] using the *RUVr* function for k = 2 factors of unwanted variation. Subsequently, differential expression analysis was performed using the negative binomial distribution-based method implemented in DESeq2[5] on the normalized summed exon read counts per gene. Pair wise comparisons between patients and HC groups were tested using the Wald test, and genes with a nominal p-value <0.05 were considered to be significantly differential. Gene expression levels are given as variance stabilised data (VSD), calculated according to DESeq2 instructions.

**Weighted gene co-expression network analysis**

Weighted gene co-expression networks were constructed using the Bioconductor/R package WGCNA[6], using the VSD data of all genes with a normalized read count expression higher than 5.53 (translates to at least 1 raw count) in all samples as input. We used a soft threshold power of 5 to construct an unsigned network with scale free topology. Modules were identified using the *cutreeDynamic* function with a minimum module size of 50 genes. Next, closely related modules were merged using the *mergeCloseModules* function, with a cutHeight of 0.25, according to the WGCNA manual.

**cDC2 cultures**

For functional experiments, cDC2s were purified from healthy donor buffy coat blood samples (Sanquin, Amsterdam, The Netherlands) as described above. Freshly isolated cDC2s were cultured in RPMI 1640 medium with GlutaMAX™ (Life Technologies), supplemented with 10% heat inactivated fetal bovine serum (Biowest) and 1% penicillin streptomycin (Life Technologies). For stimulation experiments, cDC2s were cultured at a concentration of 0.5 × 10^5^ cells/mL in 100µl in a 96-well round-bottom plate. Cells were either left untreated, or treated for 18 hours with one of the following stimuli: TLR7/8 ligand Resiquimod ( R848, 100ng/mL, Invivogen), GM-CSF (800U/mL, R&D), CXCL4 (10ug/mL, Peprotech), TNFα (100ng/mL, Tebu-bio), IFNα-2a (1000U/mL, Cell Sciences), LPS EB Ultrapure (100ng/mL, Invivogen), IL-6, (50ng/mL, ImmunoTools), IL-15 (200 ng/mL, ImmunoTools) or TGFβ-b2 (100ng/ml, R&D). For hypoxia experiments, cells were cultured in atmospheric or hypoxic conditions (Rasquinn invivO2 1000 hypoxic chamber, set at 1% O_2_ and 5% CO_2_) for 24 hours. For experiments using NR4A agonists, cells were pre-treated for one hour with either dimethyl sulfoxide (DMSO, Sigma) or NR4A agonists C-DIM5 (Tocris Bioscience) or C-DIM12 (Tebu-Bio) prior to stimulation. cDC2 cultures were incubated at 37°C in the presence of 5% CO2, for the time points indicated in each single experiment in the results section. Supernatants were stored at -80°C until further use, while cDC2s were lysed in RLTplus buffer and stored at -20°C until further use.

**PBMC cultures**

For functional experiments, three batches (individual days) of randomly selected dcSSc and matched healthy control samples of liquid nitrogen stored PBMCs were thawed in RPMI 1640 (Thermo Fisher Scientific) supplemented with 20% FCS (Sigma), and washed with PBS. Cells were resuspended in RPMI 1640 medium with GlutaMAX™ (Life Technologies), supplemented with 10% heat inactivated fetal bovine serum (Biowest) and 1% penicillin streptomycin (Life Technologies), and plated at a concentration of 0,75×10^6^ cells in 200µl in round-bottom 96-wells plates. PBMCs were pre-treated for one hour with 10uM dimethyl sulfoxide (DMSO, Sigma), C-DIM5 (Tocris Bioscience) or C-DIM12 (Tebu-Bio) prior to stimulation with R848 (100ng/mL, Invivogen) and GolgiStop (1500x, BD biosciences), and incubated at 37°C in the presence of 5% CO2, for 4 hours. IL-6 production by cDC2s was measured using intracellular cytokine staining by FACS.

**Reverse Transcription Quantitative Real-Time PCR (RT-qPCR)**

Purified RNA was reverse transcribed using the SuperScript® IV Reverse Transcriptase kit (Invitrogen), according to the manufacturer’s instructions. Gene expression was quantified, in duplicate, by RT-qPCR using the SYBR Select Master Mix (Applied Biosystems), using gene-specific primers (given below) on the QuantStudio 12k flex System (Applied Biosystems). Relative gene expression was determined according to the comparative CT (ΔΔCT) method using GUSB as an endogenous control (where the ΔCT equals the CT of the mRNA of interest—the CT of GUSB). The fold change (FC) of each sample was calculated in relation to the ΔCt of the medium control according to the formula FC = 2^−ΔΔCt^, where ΔΔCt = ΔCt sample—ΔCt reference.

**Gene specific primers used for RT-qPCR**

| **Gene** | **Forward** | **Reverse** |
| --- | --- | --- |
| GUSB | CACCAGGGACCATCCAATACC | GCAGTCCAGCGTAGTTGAAAAA |
| NR4A1 | ACTGCCCTGTGGACAAGAG | CTGTTCGGACAACTTCCTTC |
| NR4A2 | GGACTCCCCATTGCTTTTC | AGGCGAGGACCCATACTG |
| NR4A3 | CTGCCCAGTAGACAAGAGAC | CTCCTCCCTTTCAGACTATC |
| IL-6 | GGCACTGGCAGAAAACAACC | GCAAGTCTCCTCATTGAATCC |

**Assessment of IL-6 production using ELISA**

Concentrations of IL-6 in cell-free supernatants from cultured cDC2s were measured by sandwich enzyme linked immunosorbent assay (ELISA). IL-6 was quantified using the PeliKine compact human IL-6 (Sanquin Reagents, Amsterdam, The Netherlands), according to the manufacturer’s instructions.

**cDC2 / CD4+ T-cell co-cultures**

cDC2s and CD4+ T-cells were isolated in parallel from PBMCs obtained from healthy donor buffy coats using the MACS human CD1c dendritic cell isolation kit and CD4+ T Cell Isolation Kit (Miltenyi Biotec) on the autoMACS Pro Separator (Miltenyi Biotec) according to the manufacturer’s instructions. cDC2s and T-cells were cultured, in parallel, in culture medium (RPMI 1640 medium with GlutaMAX™ (Life Technologies), supplemented with 10% heat inactived fetal bovine serum (Biowest) and 1% penicillin streptomycin (Life Technologies)). cDC2s were pre-treated for one hour with either dimethyl sulfoxide (DMSO, Sigma) or specific NR4A agonists C-DIM5 (Tocris Bioscience) or C-DIM12 (Tebu-Bio), and were either left untreated or treated with R848 (100ng/mL, Invivogen). After overnight incubation, cDC2s were washed twice with sterile PBS, resuspended in culture medium and added to the CD4+ T-cells in a 1:5 ratio (10,000 cDC2s : 50,000 CD4+ T-cells, in 100µl in a round bottom 96-wells plate). Cells were cultured for 3 days at 37°C in the presence of 5% CO_2_. After 3 days, co-cultures were re-stimulated with phorbol myristate acetate (PMA, 50ng/ml, Sigma-Aldrich) and ionomycin (500ng/ml, Sigma-Aldrich) for 6 hours. During the last 3 hours of incubation, GolgiStop (1500x, BD biosciences) was added to block cytokine release. IFNy production by CD4+ T-cells was measured using intracellular cytokine staining by flow cytometry.

**Flow cytometry**

For cDC2/T-cell co-cultures and PBMC cultures, cells were washed with cold PBS and incubated with Fixable Viability Dye eF780 (eBioscience) at room temperature for 10 minutes. Cells were then transferred to V‐bottomed plates (Greiner Bio‐one), washed with PBS and incubated for 30 minutes at 4˚C in the dark with the surface staining antibodies provided below. Next, the cells were washed in FACS buffer (1% bovine serum albumin and 0.1% sodium azide in phosphate buffered saline), and fixed/permeabilized for 30 minutes at 4˚C in the dark with 100 µl Fixation/Permeabilization Concentrate and Diluent (Cat #00-5123-43, #00-5223-56, eBioscience), followed by intracellular staining using the antibodies provided below. After 60 minutes of staining at 4˚C in the dark, the cells were washed and taken up in FACS buffer and flow cytometric analyses were performed on the BD LSRFortessa with four lasers (405, 488, 561, and 635 nm) using FACSDiva software version 8.0.1. Analysis of FCS files were performed using FlowJo software (TreeStar inc.).

**Antibody panels for used for flow cytometry**

|  | **Antibody** | **Label** | **Clone** | **Company** |
| --- | --- | --- | --- | --- |
| PBMC panel | Fixable viability | eF780 | N.A. | eBioscience |
|  | Surface CD11c | PerCP-Cy5.5 | 3.9 | Thermo Scientific |
|  | Surface CD1c | APC | AD5-8E7 | Miltenyi |
|  | Surface CD19 | AF700 | HIB19 | eBioscience |
|  | Surface CD56 | AF700 | B159 | BD |
|  | Surface CD3 | PE/Cy7 | UCHT1 | Biolegend |
|  | Surface CD4 | BV510 | RPA-T4 | Biolegend |
|  | Surface HLA-DR | BV605 | G46-6 | BD |
|  | Surface CD14 | BV785 | M5E2 | Biolegend |
|  | Intracellular IL-6 | PE | MQ2-6A3 | BD |
| Co-culture panel | Fixable viability | eF780 | N.A. | eBioscience |
|  | Surface CD3 | AF700 | UCHT1 | Biolegend |
|  | Surface CD4 | AF488 | RPA-T4 | BD |
|  | Surface CD1c | APC | AD5-8E7 | Miltenyi |
|  | Intracellular IFNy | PE | 4S.B3 | BD |

**Chromatin immunoprecipitation sequencing (ChIP-seq)**

cDC2s were isolated from PBMCs obtained from healthy donor buffy coats using the MACS human CD1c dendritic cell isolation kit. 1.5×10^6^ freshly isolated cDC2s were cultured overnight in RPMI 1640 medium with GlutaMAX™ (Life Technologies), supplemented with 10% heat inactivated fetal bovine serum (Biowest) and 1% penicillin streptomycin (Life Technologies), and were either left untreated or treated with TLR7/8 agonist R848 (Resiquimod, 100ng/mL, Invivogen). After culture, cDC2s were washed with PBS and crosslinked using the truChIP® Ultra-Low Chromatin Shearing Kit (Covaris), according to the manufacturer’s instructions. Chromatin shearing was performed using AFA Fiber Pre-Slit Snap-Cap microTUBEs (Covaris) by sonication (peak incident power: 105, duty factor: 2%, cycles per burst: 200, treatment time: 12 minutes) on the Covaris S220 focused ultrasonicator (Woburn, MA, USA). After shearing, the chromatin was transferred into pre-chilled microcentrifuge tubes and centrifuged at 10,000 x g, 4˚C for 5 minutes to pellet insoluble material, and the chromatin was stored at -20˚C for downstream processing. For every condition we obtained three biological replicates, for which material from three to four donors was pooled to obtain enough chromatin for chromatin immunoprecipitation. Chromatin immunoprecipitation was performed with 3µg anti-NR4A1 (NB100-56745, Novusbio), anti-NR4A2 (NB110-40415, Novusbio) or anti-NR4A3 (NLS2341, Novusbio) antibodies, using the low cell ChIP-seq kit (Active Motif), according to the manufacturer’s instructions. For all conditions, 10% of the input chromatin was removed prior to addition of the antibodies and used to normalize the amount of immunoprecipitated DNA (input control). Following immunoprecipitation, ChIP DNA was de-crosslinked in the presence of NaCl and Proteinase K (Active Motif) in a thermocycler at 65°C overnight, and DNA was extracted by penol:chloroform:isoamylalcohol precipitation and dissolved in Low-EDTA TE buffer (Active Motif). ChIP-seq libraries were generated by GenomeScan (Leiden, the Netherlands) with the NEBNext® Ultra II DNA Library Prep kit (Illumina), and were sequenced using Illumina NovaSeq6000 generating ~20 million 150bp paired ended reads for each sample.

**ChIP-seq analysis**

Quality check of the raw sequencing reads was performed using the FastQC tool. Sequencing reads were mapped against the reference genome GRCh38.p13 built from the human genome (NCBI) using bowtie2[7]. Peaks were called using MACS2[8], by comparing the IP samples to their matched input samples in paired end mode (BAMPE). After calling, peaks from ENCODE blacklist regions[9], and X- and Y-chromosome peaks were filtered out to reduce noise and exclude sex-specific peaks. Peaks were annotated to the nearest genes using the ChIPseeker Bioconductor/R package[10]. Peaks were considered to be associated to a gene when they were annotated within a 10kb range (up- or downstream) to the gene transcription start site (TSS).

**Statistical analysis**

The Mann Whitney test was used to compare any combination of two groups. For multiple group comparisons, the one- or two-way analysis of variance (ANOVA) was used. A *p*-value < 0.05 was considered statistically significant. Figures were produced using the R package *ggplot2*[11]. GO-term and KEGG enrichment analyses were performed using enrichGo and enrichKEGG functions from the clusterProfiler R package[12]. Terms with a Benjamini and Hochberg (B&H) corrected *p*-value <0.05 were considered significant. Spearman's rank correlation coefficient was calculated to assess correlations.

1. A. Dobin et al., “STAR: ultrafast universal RNA-seq aligner,” Bioinformatics, vol. 29, no. 1, pp. 15–21, Jan. 2013
2. S. Anders, P. T. Pyl, and W. Huber, “HTSeq--a Python framework to work with high-throughput sequencing data.,” Bioinformatics, vol. 31, no. 2, pp. 166–169, Jan. 2015
3. D. Risso, K. Schwartz, G. Sherlock, and S. Dudoit, “GC-Content Normalization for RNA-Seq Data,” BMC Bioinformatics, vol. 12, no. 1, p. 480, 2011
4. D. Risso, J. Ngai, T. P. Speed, and S. Dudoit, “Normalization of RNA-seq data using factor analysis of control genes or samples,” Nat. Biotechnol., vol. 32, no. 9, pp. 896–902, Sep. 2014
5. M. I. Love, W. Huber, and S. Anders, “Moderated estimation of fold change and dispersion for RNA-seq data with DESeq2,” Genome Biol., vol. 15, no. 12, p. 550, Dec. 2014
6. P. Langfelder and S. Horvath, “WGCNA: an R package for weighted correlation network analysis.,” BMC Bioinformatics, vol. 9, no. 1, p. 559, Dec. 2008
7. B. Langmead and S. L. Salzberg, “Fast gapped-read alignment with Bowtie 2,” Nat. Methods, vol. 9, no. 4, pp. 357–359, Apr. 2012
8. J. M. Gaspar, “Improved peak-calling with MACS2,” bioRxiv. 2018
9. H. M. Amemiya, A. Kundaje, and A. P. Boyle, “The ENCODE Blacklist: Identification of Problematic Regions of the Genome,” Sci. Rep., vol. 9, no. 1, p. 9354, Dec. 2019
10. G. Yu, L.-G. Wang, and Q.-Y. He, “ChIPseeker: an R/Bioconductor package for ChIP peak annotation, comparison and visualization,” Bioinformatics, vol. 31, no. 14, pp. 2382–2383, Jul. 2015
11. H. Wickham, ggplot2: Elegant Graphics for Data Analysis. Springer-Verlag New York, 2016.
12. G. Yu, L.-G. Wang, Y. Han, and Q.-Y. He, “clusterProfiler: an R Package for Comparing Biological Themes Among Gene Clusters,” Omi. A J. Integr. Biol., vol. 16, no. 5, pp. 284–287, May 2012
