## Supplementary Figures for "Nuclear receptor subfamily 4A signaling as a key disease pathway of CD1c+ dendritic cell dysregulation in systemic sclerosis"

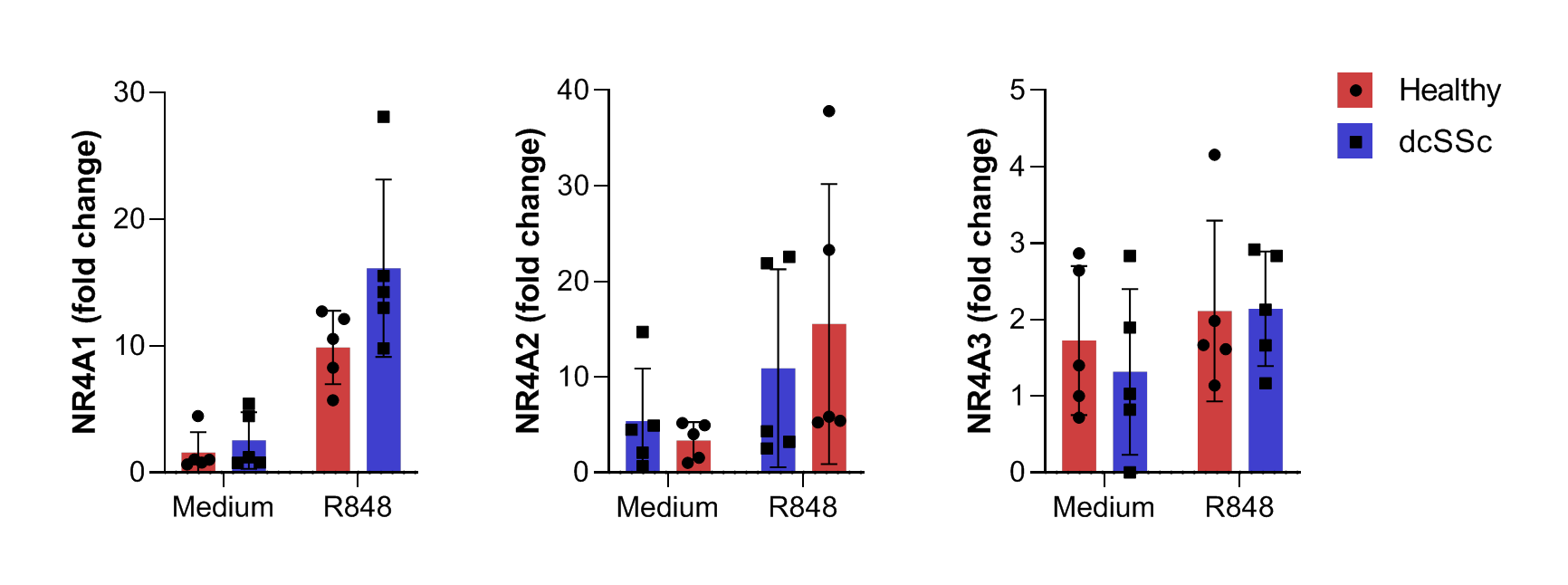

**Supplementary Figure 1: RT-qPCR of NR4A expression in stimulated CD1c+ cDCs from healthy controls and dcSSc patients.** NR4A1, NR4A2 and NR4A3 for untreated (medium) or treated (R848) cDCs following 18 hours of culture. Relative mRNA expression levels (FC) shown are normalized to GUSB housekeeping levels. Bars depict mean, error bars depict SEM.

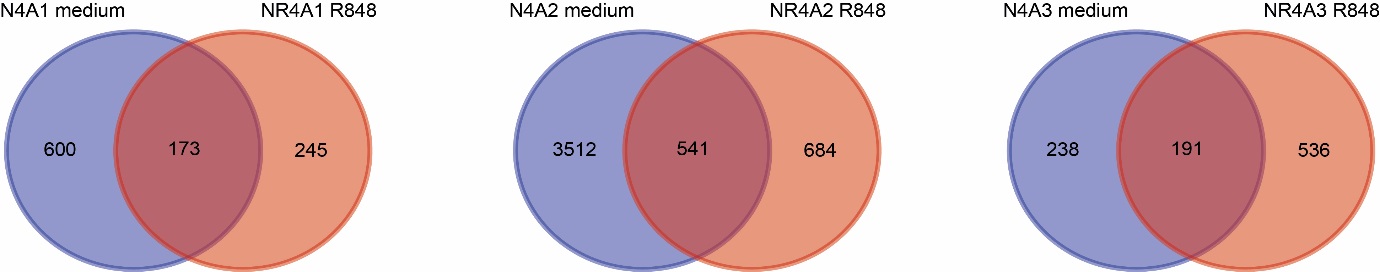

**Supplementary Figure 2. Genome-wide identification of NR4A binding sites in resting and activated CD1c + cDCs.** Venn diagrams depicting the overlap of NR4A1, NR4A2 and NR4A3 transcription factor binding sites (within 10kb of the nearest gene promoter region) in resting (blue, medium) and activated (red, R848) cDCs.

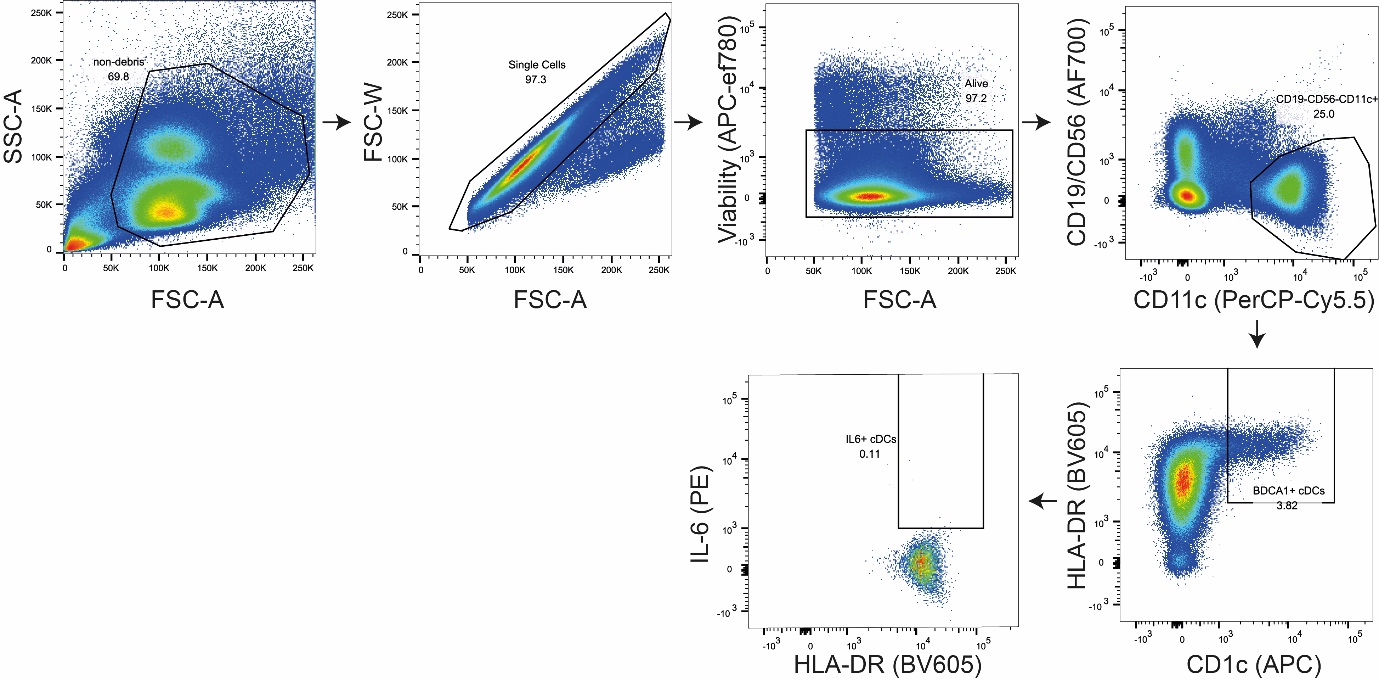

**Supplementary Figure 3: Gating strategy for measuring IL-6 production in the CD1c+ cDC fraction in PBMC cultures.**

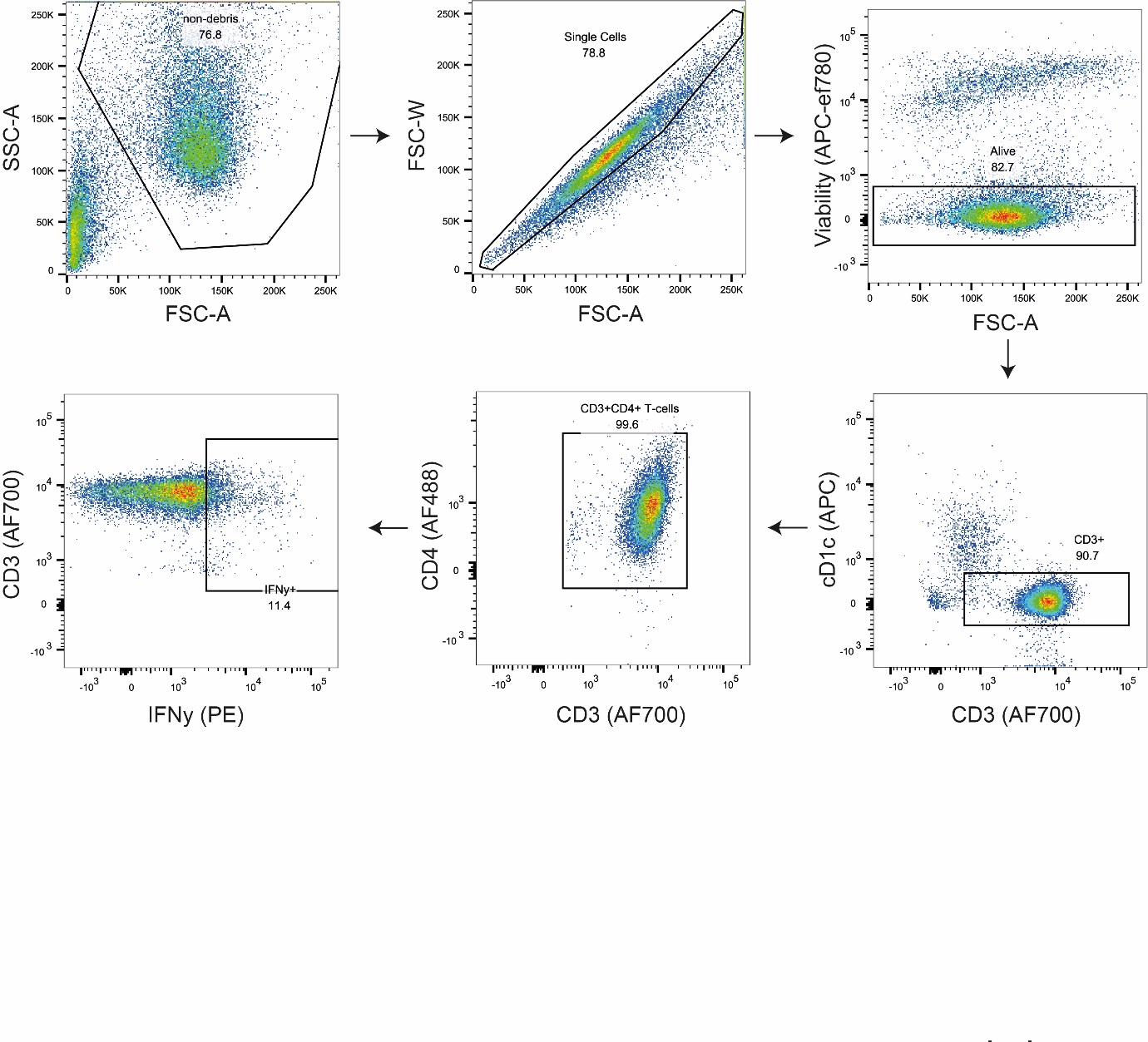

**Supplementary Figure 4: Gating strategy for measuring IFNy production by CD4+ T-cells in cDC/T-cell co-cultures.**
